# Loss of PARP7 increases type I interferon signalling and prevents pancreatic tumour growth by enhancing immune cell infiltration

**DOI:** 10.1101/2024.09.18.613621

**Authors:** Vinicius Kannen, Marit Rasmussen, Siddhartha Das, Paolo Giuliana, Fauzia N. Izzati, Hani Choksi, Linnea A. M. Erlingsson, Ninni E. Olafsen, Paola Cappello, Indrek Teino, Toivo Maimets, Kristaps Jaudzems, Antanas Gulbinas, Zilvinas Dambrauskas, Landon Edgar, Denis M. Grant, Jason Matthews

## Abstract

Pancreatic ductal adenocarcinoma (PDAC) is one of the most lethal forms of cancer, and despite low incidence rates, it remains the sixth leading cause of cancer related deaths worldwide. Immunotherapy, which aims to enhance the immune system’s ability to recognize and eliminate cancer cells, has emerged as a promising approach in the battle against PDAC. PARP7, a mono-ADP-ribosyltransferase, is a negative regulator of the type I interferon (IFN-I) pathway and has been reported to reduce anti-tumour immunity. Using murine pancreatic cancer cells, we found that loss of *Parp7* elevated the levels of interferon stimulated gene factor 3 (ISGF3) and its downstream target genes, even in the absence of STING. Cancer cells deficient in *Parp7* produced smaller tumours when injected into immunocompetent mice. Transcriptomic analyses revealed that tumours knocked out for *Parp7* (Parp7^KO^) had increased expression of genes involved in immunoregulatory interactions and interferon signalling pathways. Characterization of tumour infiltrating leukocyte (TIL) populations showed that Parp7^KO^ tumours had higher proportions of natural killer cells, CD8 T cells and a lower proportion of anti-inflammatory macrophages (M2). The overall TIL profile of Parp7^KO^ tumours was suggestive of a less suppressive microenvironment. Our data show that loss of *Parp7* reduces PDAC tumour growth by increasing the infiltration of immune cells and enhancing anti-tumour immunity. These findings provide support to pursue PARP7 as a therapeutic target for PDAC.

## Introduction

Pancreatic ductal adenocarcinoma (PDAC), which represents more than 90% of pancreatic cancers (PC), is a highly aggressive disease and poses a challenge in the field of oncology [1]. The disease arises from the uncontrolled growth of malignant cells in the pancreas, an essential organ responsible for producing digestive enzymes and hormones, including insulin. Despite low incidence rates, PDAC remains the sixth leading cause of cancer-related deaths worldwide [2]. Due to its anatomical location, PDAC is characterized by its silent progression, often remaining undetected until advanced stages, resulting in poor prognoses and high mortality rates [3]. Although there have been significant advancements in PDAC research, the mortality to incidence ratio has changed little over the past few decades. Compared with other cancers, PDAC exhibits remarkable resistance to conventional therapies and possesses a highly immunosuppressive tumour microenvironment (TME), enabling cancer cells to “hide” from the immune system [4]. Surgery with curative intent together with adjuvant chemotherapy is the treatment of choice; however, this is only possible in about 10-20% of patients [5]. Recent progress in targeting the immune system for cancer treatment, referred to as cancer immunotherapy, has caused a paradigm shift in therapeutic options for cancer patients. One of the most studied strategies involves targeting the immunosuppressive interaction between programmed death ligand 1 (PD-L1), which is present on tumour cells, and its receptor, programmed death receptor (PD-1), which is expressed on immune cells, such as activated T cells, natural killer (NK) cells, B cells, macrophages and different subsets of dendritic cells (DCs) [6]. Inhibition of the PD-1/PD-L1 immune checkpoint axis has produced impressive response rates in various malignancies, such as melanoma, renal and lung cancer. However, despite PD-L1 being expressed in human PC samples, immunotherapy targeting PDAC has so far been ineffective due to reduced immune cell infiltration [7, 8]. Thus, augmented engagement of the immune response may provide significant clinical benefit for PDAC patients.

Type I Interferons (IFN-Is) are cytokines that are released in response to pathogen or damage associated molecular patterns (PAMPs or DAMPs, respectively). They are expressed by almost all cells in the body and are involved in the regulation of many biological processes, such as cellular immune responses to infections, cell cycle regulation, differentiation and apoptosis [9]. Nucleic acids released from damaged cells function as DAMPs and can be recognized by pattern recognition receptors (PRRs), thereby eliciting an immune response. The presence of cytosolic DNA activates cyclic GMP-AMP synthase (cGAS), which synthesizes cyclic guanosine monophosphate-adenosine monophosphate (cGAMP). cGAMP subsequently activates stimulator of interferon response cGAMP interactor (STING) resulting in activation of TANK binding kinase 1 (TBK1), which phosphorylates IFN regulatory factor 3 (IRF3). IRF3 homodimerizes, translocates to the nucleus, and upregulates expression of IFN-Is, such as IFNβ, which is secreted from the cell [10]. IFNβ binds to the IFNα/β receptor (IFNAR) complex on immune and non-immune cells, resulting in activation of signal transducer and activator of transcription 1 (STAT1) and STAT2. These proteins in turn form a complex with interferon regulatory factor 9 (IRF9) known as IFN stimulated gene factor 3 (ISGF3), which regulates the expression of IFN stimulated genes (ISGs) that play important roles in immunity [11]. STING has emerged as a promising target for cancer therapy, providing new strategies to exploit the immune system to combat cancer [12]. The expression levels of IFN-Is and other inflammatory cytokines downstream of STING activation contribute to enhanced anti-tumour immune responses [13]. By utilizing this mechanism, STING agonists have been shown to induce tumour regression by enhancing the ability of immune cells to target cancer cells [14]. IFN-Is regulate tumour infiltrating immune cells and are critically important in maintaining antigen-presenting lymphocyte function for effective anti-tumour immunity. They also exhibit cancer cell intrinsic properties, such as growth inhibition and increased apoptosis [11]. Increasing IFN-I levels has shown promising results in the preclinical and clinical setting, and some studies report that they synergize with immune checkpoint inhibitors to reduce tumour growth in cancer models [15–18]. In contrast, loss of IFN-I signalling enhances tumorigenesis and impairs anti-tumour responses [19].

PARP7, also known as 2,3,7,8-tetrachlorodibenzo-*p*-dioxin (TCDD)-inducible poly-ADP-ribose polymerase (TIPARP) is a member of the ADP-ribosyltransferase diphtheria-like (ARTD) family and a critical regulator of innate immune signalling [20]. PARP7 uses NAD^+^ to transfer one molecule of ADP-ribose to specific amino acid residues on itself and on target proteins, in a process referred to as mono-ADP-ribosylation (MARylation) [21]. MARylation is a reversible post-translational modification involved in several biological processes, such as immune cell function, transcriptional regulation, and DNA repair [22]. PARP7 mRNA expression is regulated by several transcription factors, including the ligand-induced transcription factor aryl hydrocarbon receptor (AHR). In turn, PARP7 acts as a negative regulator of AHR via MARylation [23, 24]. Previous studies have reported that PARP7 is required for AHR-dependent repression of IFN-I responses during viral infection associated with PAMP stimulation, a process that requires its catalytic activity [25]. The repressive actions of PARP7 have been attributed to its ability to MARylate TBK1, thus preventing the downstream upregulation of IFN-Is [25]. More recent studies provide evidence that PARP7 regulates IFN-I signalling downstream of TBK1 [26] and by targeting nuclear factor kappa B [27].

Treatment with IFN-I inducers in combination with immune checkpoint inhibitors has been shown to sustain anti-tumour responses in models of aggressive cancers [10]. Recent studies in preclinical mouse models, showed that pharmacological inhibition of PARP7 with RBN-2397 reduced CT26 colon tumour growth in immunocompetent mice, which was dependent on IFN-I signalling and that cotreatment of RBN-2397 with anti-PD1 further reduced tumour growth compared with either treatment alone [28]. Consistent with these findings, we reported that injection of mice with murine EO771 breast cancer cells in which PARP7 was knocked out (Parp7^KO^) resulted in >80% reduced tumour growth in *Parp7* deficient mice compared with injected wildtype EO771 cells in wildtype (WT) mice [27]. This was due to an increased infiltration of tumour-associated immune cells, resulting in augmented anti-tumour immunity. These findings show that PARP7 loss or its inhibition reduces tumour growth in different preclinical models by increasing anti-tumour responses. However, PARP7’s involvement in PDAC remains unclear.

In this study, we show that loss of PARP7 expression or its activity increases basal ISG expression levels in murine pancreatic cancer cells *in vitro*, and we demonstrate that *Parp7* loss decreases tumour growth through increased tumour infiltrating immune cells and enhanced anti-tumour immunity. Our results suggest that targeting PARP7 alone or combination with other immunotherapies should be considered as a new therapeutic strategy against PDAC.

## Methods

### Chemicals and plasmids

DMSO was purchased from Sigma Aldrich (St. Louis, MO, USA), DMXAA from Invivogen (San Diego, CA, USA), RBN-2397 from MedChemExpress (Monmouth Junction, NJ, USA), FICZ from SelleckChem (Houston, TX, USA), and IFNβ from R&D Systems (Minneapolis, MN, USA). The pSpCas9(BB)-2A-Puro (PX459) plasmid was purchased from Addgene (plasmid #62988) (Watertown, MA, USA).

### Cell culture

CR705 cells derived from a spontaneous pancreatic tumour in a *LSL-Kras^G12D/+^; LSL-Trp53^R172H/+^; Pdx1-Cre* (KPC) mouse were used in this study [29]. Cells were maintained in RPMI culture medium (1.0 g/L glucose), supplemented with 10% *v/v* heat-inactivated fetal bovine serum (FBS), 1% *v/v* L-glutamine and 1% *v/v* penicillin-streptomycin. Cells were cultured at 37°C with 100% humidity and 5% CO_2_, and subcultured when confluency reached 80%.

### Generation of PARP7 deficient cells

The guide oligos used to make the gRNA were: 5’-CACCGTCTTCTCAGAAATTCTCATT-3’ and 5’-AAACAATGAGAATTTCTGAGAAGAC-3’. After inserting the resulting gRNA into the PX459 plasmid, the cells were transfected, selected and expanded as previously described [27]. To confirm knockout, genomic DNA from several clones was harvested, and the target site was amplified and sequenced. The primers used for sequencing were: Forward 5’-TGCAGATTTTTGCATAGCTTTTG-3’ and reverse 5’-TTGTCTTGGAAAGCTC CTGGT-3’. After screening, one clone was subsequently expanded, and further analysed. Cells transfected with an empty PX459 plasmid were also selected and expanded and will be referred to as CR705 WT cells.

### Western blotting

Cells used for western blotting were seeded in six-well plates and treated the following day. Cells were lysed in TE-buffer supplemented with 1% *w/v* SDS. After brief sonication, the samples were boiled at 95°C for 10 min. Protein concentration was determined with a BCA assay (Thermo Fisher Scientific, Waltham, MA, USA). Proteins were separated by SDS-PAGE and transferred to polyvinylidene fluoride (PVDF) membranes. The antibodies used were: lab generated anti-PARP7 antibody [30], anti-AHR (Enzo Life Sciences, Farmingdale, NY, USA; bml-sa210-0100), anti-STING (Cell Signalling Technology, Danvers, MA, USA; D2P2F), anti-STAT1 (Cell Signalling Technology; #9172), anti-STAT2 (Cell Signalling Technology; D9J7L), anti-IRF9 (Cell Signalling Technology; D9I5H), anti-pSTAT1 (Y701) (Cell Signalling Technology; D4A7), and anti-β-actin (Sigma-Aldrich; AC-74). After incubation with corresponding secondary antibody (Rabbit, Mouse, Cell Signalling Technology), the protein bands were visualized with SuperSignal™ West Dura Extended Duration Substrate or SuperSignal™ West Atto Ultimate Sensitivity Substrate (Thermo Fisher Scientific, Waltham, MA, USA).

### Real time qPCR (RT-qPCR)

Total RNA was isolated using the Aurum™ Total RNA isolation kit (BioRad, Hercules, CA, USA), and used to synthesize cDNA with the High-Capacity cDNA Reverse Transcription Kit (Applied Biosystems, Waltham, MA, USA). The RT-qPCR assays were set up as previously described [30]. The primers used are provided in **Supplementary Table S1**.

### Proliferation assays

Cells were seeded in 96-well plates on day 0 at a density of 2000 cells per well. On day 1, CellTiter Glo (Promega, Madison, WI, USA) assay was performed according to the manufacturer’s instructions to obtain baseline measurements. The remaining cells were treated with DMSO or 100 nM RBN-2397. On day 4, the treated cells were measured. Data are shown as a relative increase in proliferation compared to the baseline measurements.

### Mouse models and tumour studies

Female immune deficient (NOD.Cg-*Prkdc^scid^ Il2rg^tm1Wjl^/SzJ,* NSG: #005557) and immunocompetent (C57BL/6J: #000664) mice were purchased from The Jackson Laboratory (Bar Harbor, ME, USA). Mice aged 8-12 weeks underwent isoflurane anaesthesia before a single subcutaneous injection with either CR705 WT or Parp7^KO^ cells. CR705 cells were prepared as single cell suspensions and injected at a density of 5 × 10^5^ cells. Tumour growth was monitored with caliper measurements, and tumour volumes were calculated using the standard formula π/6 × W^2^ × L. Mice were euthanized at the end of experiments by cervical dislocation, and tumours were prepared for histological analyses. All experimental animals were housed in the Division of Comparative Medicine at the University of Toronto, with a 12 h light/dark cycle, and access to chow and water *ad libitum*. Care and treatment of animals followed the guidelines set by the Canadian Council on Animal Care and was approved by the University of Toronto Animal Care Committee.

### Immunohistochemistry

Sectioning and staining of the tumours were performed according to standard methods. Fixed tissues were provided to the HistoCore Facility at the Princess Margaret Cancer Centre (Toronto, Ontario, Canada), and sample processing, staining with Ki67, CD3, and CD8 and scanning were done at the facility. Quantification analysis was performed with QuPath v0.4.3 [31].

### RNA sequencing and data analysis

Total RNA was isolated from approximately 100 mg of tumours from CR705 WT or Parp7^KO^ cells using the Aurum™ Total RNA isolation kit (BioRad) according to the manufacturer’s protocol. The raw RNA sequence paired-end fastq files were quantified using the Salmon tool with “libtype” flag as automatic and mm10 version of the Salmon index file [32]. The index was generated using the salmon “index” flag with the mm10 transcripts fasta file supplied. The “tximport” import function from the tximport package (v1.26.1; [33]) was used to import the Salmon quantification data for further processing including differential expression analysis by DESeq2 [34]. For all comparisons, WT tumour samples were considered as the control. Significant genes were considered as those with absolute log fold change greater than 1 and Benjamin Hochberg false discovery rate value of differential expression less than 0.01 and tested using the Wald Test implemented in DESeq2. Pathway analysis was done using the Reactome database.

### Tumour and spleen dissociation into single cells

Tumours were dissected at endpoint and were processed into single-cell suspensions using a Mouse Tumour Dissociation Kit (Miltenyi Biotec, Bergisch Gladbach, Germany) according to the manufacturer’s protocol. Approximately, 1-2 mm tumour pieces were placed in an ice-cold Tissue Storage Solution (#130-100-008) before transferred into gentleMACS™ C Tubes (Miltenyi Biotec) containing the appropriate kit reagents and enzymes. The tumour samples were then incubated in a gentleMACS Octo Dissociator (Miltenyi Biotec). SmartStrainers were used to filter debris and erythrocyte lysis was performed with a Red Blood Cell Lysis Solution (Miltenyi Biotec). Cells were counted and frozen in MACS Freezing Solution and store at – 80°C.

### Spectral flow cytometer analysis

Spectral flow cytometry analysis of stained cell suspensions was performed on a Cytek Aurora spectral flow cytometer equipped with a 3-laser, 38-detector array and operated by the Cytek SpectroFlo software version 3.2.1. Healthy splenocytes or Ultra-Comp control compensation particles (Thermo Fisher Scientific) defined spectral unmixing standards data for single colour controls for each of the 28 targets analysed (**Supplementary Table S2**). Cells suspensions were washed with PBS before viability staining with Zombie NIR viability dye for 15 min at room temperature. Cells were then incubated with Fc block for 15 min on ice. Staining with Anti-CD62L, TCRγ8, and CCR6 antibodies was then performed by incubating cells with optimized antibody concentrations for 30 min at 37°C. Staining with antibodies against other cell surface receptors was then performed for 30 min at room temperature. For intracellular staining, cells were fixed and permeabilized with the FoxP3 Fix/Perm buffer set (eBioscience) before incubating with anti-FoxP3 and RORγt antibodies for 1 hour at room temperature. All staining steps with fluorescent reagents were performed in the dark. The data was analysed with FlowJo v10 software.

### Statistical analysis

Data are represented as standard error of the mean (S.E.M) of at least three individual experiments and were analysed with GraphPad Prism v8.2 (San Diego, CA, USA). Statistical analyses were done using a two-tailed student’s t-test, one-or two-way analysis of variance (ANOVA). For flow cytometry, cell counts and population frequencies were determined using FlowJo V10 software. Differences in population frequencies of different tumour infiltrating leukocyte (TIL) populations between WT and Parp7^KO^ tumours were compared using Mann-Whitney test or mixed-effects analysis with Šídák’s multiple comparisons test.

## Results

### CR705 cells express AHR and are sensitive to RBN-2397 induced decreases in cell proliferation

The CR705 cells used in this study were derived from a pancreatic tumour in the *LSL-Kras^G12D/+^; LSL-Trp53^R172H/+^; Pdx1-Cre* (KPC) mouse model [29]. This model is commonly used to study human pancreatic cancer, as the mice spontaneously develop pancreatic cancer at 11-12 weeks of age, and the tumours display many of the key features of the TME observed in human patients [35]. To characterize the effects of PARP7 inhibition or loss in these cells, we treated cells with 100 nM RBN-2397 and generated *Parp7* knockout cells using CRISPR/Cas9 gene editing. After selection and sequencing of several potential clones, only one clone contained indels that resulted in frameshift mutations in the *Parp7* gene (**Supplementary Figure S1**). This clone, referred to as Parp7^KO^, was expanded and further characterized. We have previously reported that PARP7 catalyses its own proteolytic degradation and that inhibition of PARP7 activity stabilizes its protein levels [30]. As such, treatment with the PARP7 inhibitor RBN-2397 enabled visualization of PARP7 in the WT cells, but not in the Parp7^KO^ cells, indicating successful knockout (**Figure 1A**). Because of the importance of AHR in regulating PARP7 levels, and the well-established feedback inhibition that PARP7 has on AHR signalling, we also determined AHR levels in WT and Parp7^KO^ cells. CR705 cells expressed AHR, and AHR protein levels were elevated in Parp7^KO^ cells (**Figure 1A**). When cells were treated with the endogenous AHR agonist 6-formylindolo(3,2-b)carbazole (FICZ) [36], we observed a significantly higher increase in expression levels of the AHR target gene, *cytochrome P450 1a1* (*Cyp1a1*), in the Parp7^KO^ compared with WT cells (**Figure 1B**). This is in line with previous findings showing that PARP7 acts as a negative regulator of AHR [23, 37]. We next determined whether the CR705 cells were sensitive to the antiproliferative actions of RBN-2397 or PARP7 loss. Both PARP7 inhibition and knockout significantly decreased cell proliferation (**Figure 1C**), which is consistent with AHR expression being an important factor in determining the susceptibility to the antiproliferative effects of RBN-2397 [38].

**Figure 1.**
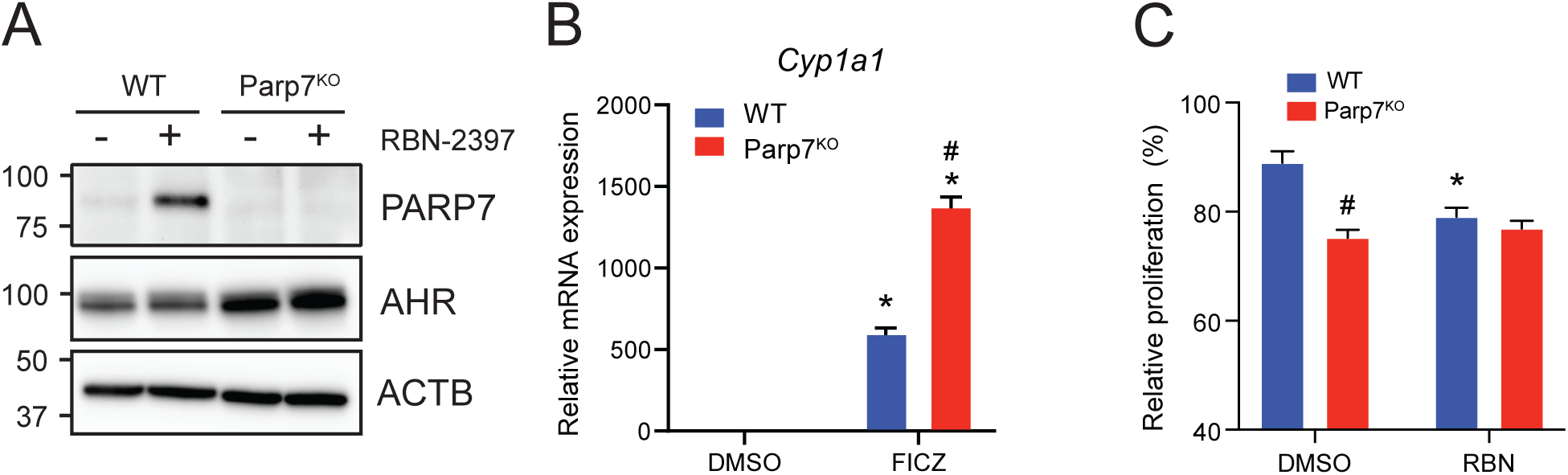

### Loss of PARP7 increases levels of ISGF3 and downstream signalling

PARP7 has been proposed as an anti-cancer therapeutic target due to its modulation of IFN-I signalling [28, 39, 40]. Because of this, we tested whether CR705 cells harboured an intact IFN-I signalling pathway. Cells were treated with RBN-2397 and co-treated with the murine specific STING agonist 5,6-dimethylxanthenone-4-acetic acid (DMXAA) (**Figure 2A**). *Ifnb1* levels were unaffected by RBN-2397 and DMXAA alone or by their co-treatment. *Ifnb1* levels did not significantly differ between the WT and Parp7^KO^ cells. Consistent with this finding, CR705 WT and Parp7^KO^ cells did not express STING (**Figure 2B**). STING expressing EO771 cells were used as positive controls. Since we previously reported that the ISGF3 complex was upregulated in PARP7 deficient cells, we tested whether this was true for CR705 cells (**Figure 2C**) [27]. As expected, we observed that PARP7 protein levels were stabilized after treatment with RBN-2397. In agreement with previous findings, the expression levels of STAT1, STAT2 and IRF9, which form ISGF3, were upregulated in the Parp7^KO^ cells, but not in the WT cells treated with RBN-2397. We were unable to detect phosphorylation of STAT1 in these samples (data not shown). In contrast to previous observations in EO771 cells, longer exposure to RBN-2397 (48 and 72 h) did not increase STAT1 levels in the WT cells (**Figure 2D**) [27]. In line with the elevated protein levels, we also observed significant increases in *Stat1* and *Irf9* mRNA levels, but not *Stat2* mRNA levels in the Parp7^KO^ cells (**Figures 2E-G**).

**Figure 2.**
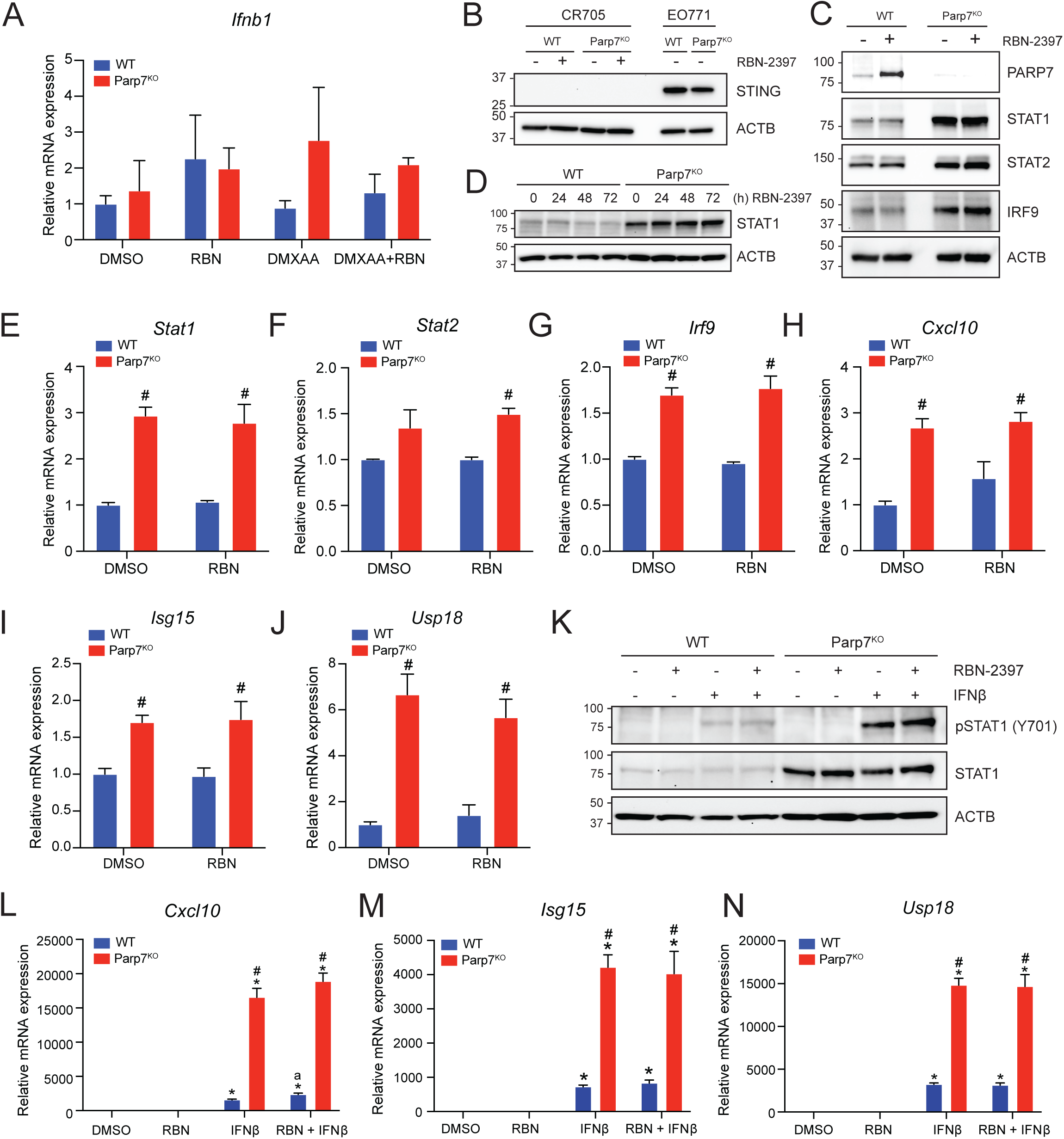

We then determined the expression levels of ISGF3 target gene *Cxcl10* to evaluate the downstream signalling pathway and found the levels to be significantly increased in Parp7^KO^ cells (**Figure 2H**) [41]. To further confirm increased ISGF3 signalling in these cells, we tested the expression level of target genes *Isg15* and *Usp18* (**Figure 2I and J**). Both *Isg15* and *Usp18* mRNA levels were significantly upregulated in the Parp7^KO^ cells, but not in WT cells treated with RBN-2397. Longer exposure to RBN-2397 (48 and 72 h) did not further elevate the expression levels of these target genes (data not shown). Exposure to exogenously added IFNβ resulted in phosphorylation of STAT1, but this was not further increased by RBN-2397 (**Figure 2K**). Parp7^KO^ cells displayed increased levels of phosphorylated STAT1, but quantification of the bands revealed that the relative levels of pSTAT1 compared to the native STAT1 were lower than in the WT cells (**Supplementary Figure 2A**). Downstream target genes *Cxcl10*, *Isg15* and *Usp18* were all significantly increased in response to IFNβ, and further elevated in the Parp7^KO^ cells (**Figure 2L-N**). Treatment with RBN-2397 slightly increased levels of *Cxcl10* in the WT cells but did not affect the levels of *Isg15* or *Usp18* (**Supplementary Figures 2B-D**) Taken together, these findings indicate that CR705 cells exhibit a partially functional IFN-I signalling pathway with increased responsiveness after *Parp7* loss. Furthermore, exposure to IFNβ revealed an intact pathway downstream of the IFNAR complex, with increased activity in the *Parp7* deficient cells. This may implicate a role for PARP7 in IFN-I signalling that is independent of upstream secretion of IFNβ.

### Loss of Parp7 affects expression of ARTD family members

Since we previously found that PARP7 loss resulted in increased expression of several members of the ARTD family [27], we determined their expression levels in CR705 WT and Parp7^KO^ cells (**Figure 3A**). Consistent with our previous findings, the levels of the MARylating enzymes *Parp9*, *Parp10* and *Parp14* were significantly upregulated in the Parp7^KO^ cells, but not in the WT cells after treatment with RBN-2397 (**Figure 3B-D**). Notably, PARP14 is also a target gene and positive regulator of the IFN-I pathway [42]. Both loss and inhibition of PARP7 resulted in decreased expression of *Parp2* and *Parp13* (Figure 3E and G), while *Parp7* loss, but not inhibition, decreased levels of *Tnks2* (**Figure 3F**). The individual graphs for the other PARPs are provided in **Supplementary Figure S3**.

**Figure 3.**
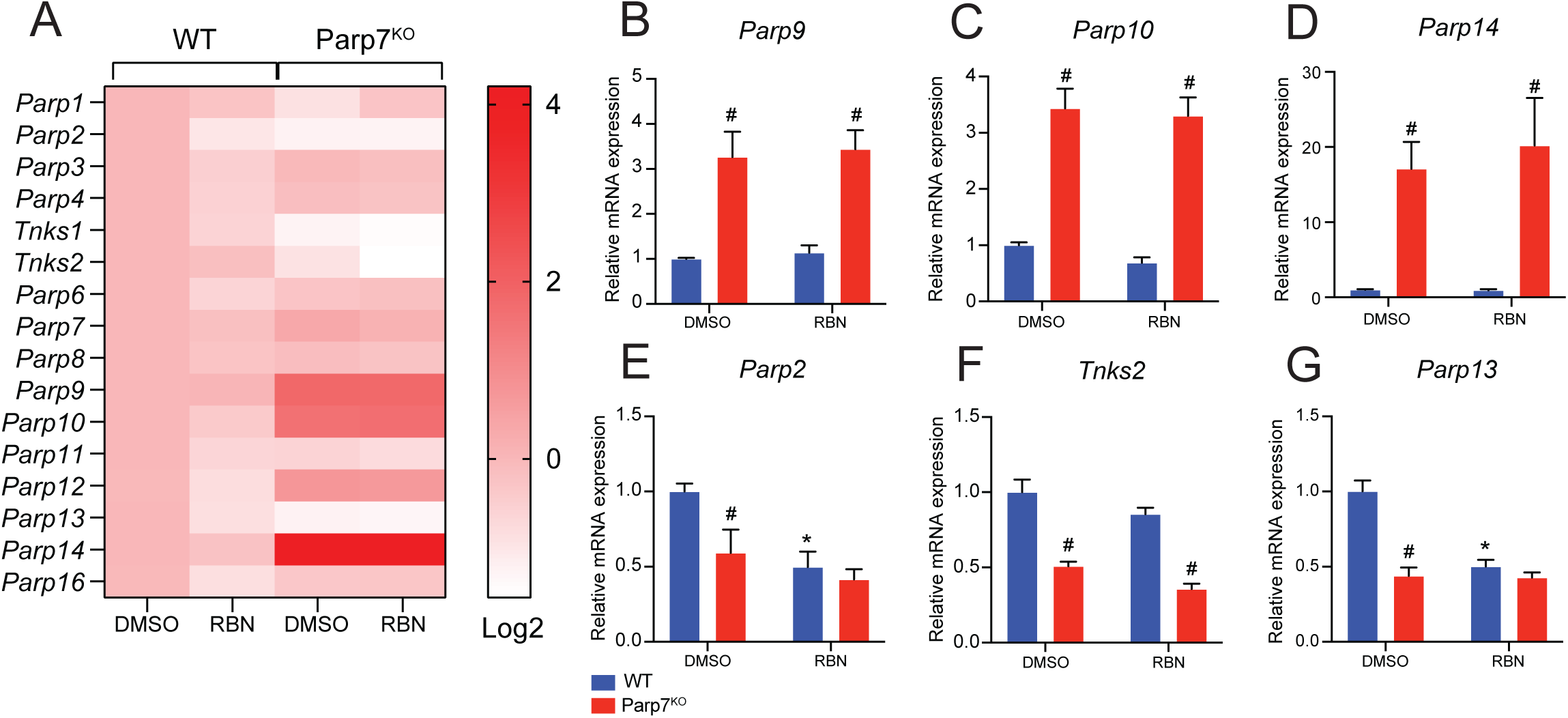

### Loss of Parp7 in CR705 PDAC cells reduces tumour growth

To determine the role of PARP7 on the ability of CR705 to form tumours *in vivo*, we injected WT and Parp7^KO^ cells into immune deficient NSG mice. Injection of Parp7^KO^ cells gave raise to tumours that grew more slowly compared with WT cells (**Figure 4A**). These findings were in line with the reduced proliferation of Parp7^KO^ cells compared with WT cells. Because IFN-I signalling was increased in Parp7^KO^ cells and it is known to enhance immune cell-mediated anti-tumour effects, we next tested the abilities of WT and Parp7^KO^ cells to form tumours in immunocompetent C57BL/6 mice. Injection of Parp7^KO^ cells gave rise to smaller tumours compared with injection of WT cells in C57BL/6 mice (**Figure 4B**) and this effect was greater than that observed in NSG mice. We did not observe any differences in the number Ki67^+^ cells, a marker of cell proliferation from WT or Parp7^KO^ tumours in C57BL/6 mice (**Figure 4C and D**). However, increased staining of T cells (CD3^+^) and specifically CD8^+^ T cells were observed in Parp7^KO^ compared with WT tumours (**Figure 4E-H**).

**Figure 4.**
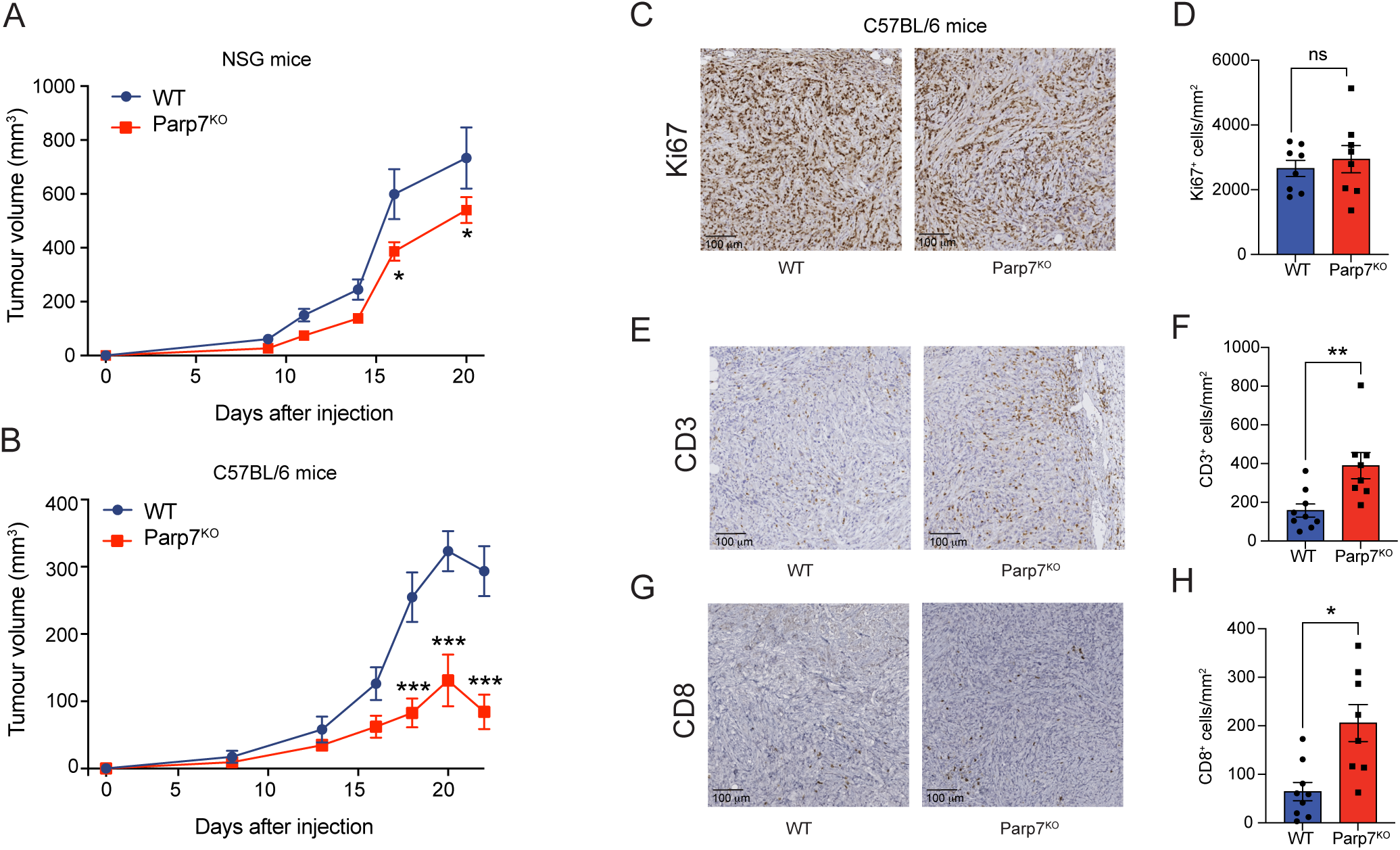

### Tumours deficient in Parp7 display a less immunosuppressive repertoire of infiltrating immune cells

Spectral flow cytometry was then used to assess differences in the proportion of tumour infiltrating leukocyte (TIL) populations between Parp7^KO^ and WT control tumours. A broad range of TILs including several lymphocyte, and myeloid lineage cell populations were identified based on their expression of characteristic cell surface markers (**Figure 5A-C, Supplementary Figure S4**). Relative proportions of each population were reported as a percentage of the total CD45^+^ leukocytes in the tumour. TILs from Parp7^KO^ tumours had a higher proportion of NK cells, eosinophils, and F4/80^High^ Sig-F^High^ Ly6G^High^ Ly6C^Low^ cells. This latter population could be inclusive of mature and long-lived neutrophils [43] and myeloid-derived suppressor cells (MDSCs) [44] (**Figure 5D and 5E**). In contrast, Parp7^KO^ tumours had a lower proportion of B cells, macrophages, and CD11b^+^ dendritic cells (CD11b^+^ DCs). Although we observed a continuum of macrophage phenotypes rather than distinct populations, we were able to use staining controls to delineate M1-like (CD80^High^ CD206^Low^) pro-inflammatory and M2-like (CD80^Low^ CD206^High^) anti-inflammatory cells (**Supplementary Figure S5A**). A similar proportion of macrophages from Parp7^KO^ and WT tumours had an M1 phenotype (**Supplementary Figure S5B**); however, a lower proportion with an M2 phenotype were observed specifically in Parp7^KO^ tumours (**Figure 5F**). The net difference in relative macrophage abundance was absorbed by populations we could not identify as either M1 or M2 (**Supplementary Figure 5A**). Further, Parp7^KO^ tumours harboured fewer F4/80^Low^ Ly-6G^High^ Ly-6C^Low^ and F4/80^High^ Ly-6G^Low^ Ly-6C^High^ cells, which have been previously defined as polymorphonuclear and monocytic MDSCs, respectively. No differences in the expression of the immune checkpoint PD-L1 were observed across all TIL populations (**Supplementary Figure S5C**).

**Figure 5.**
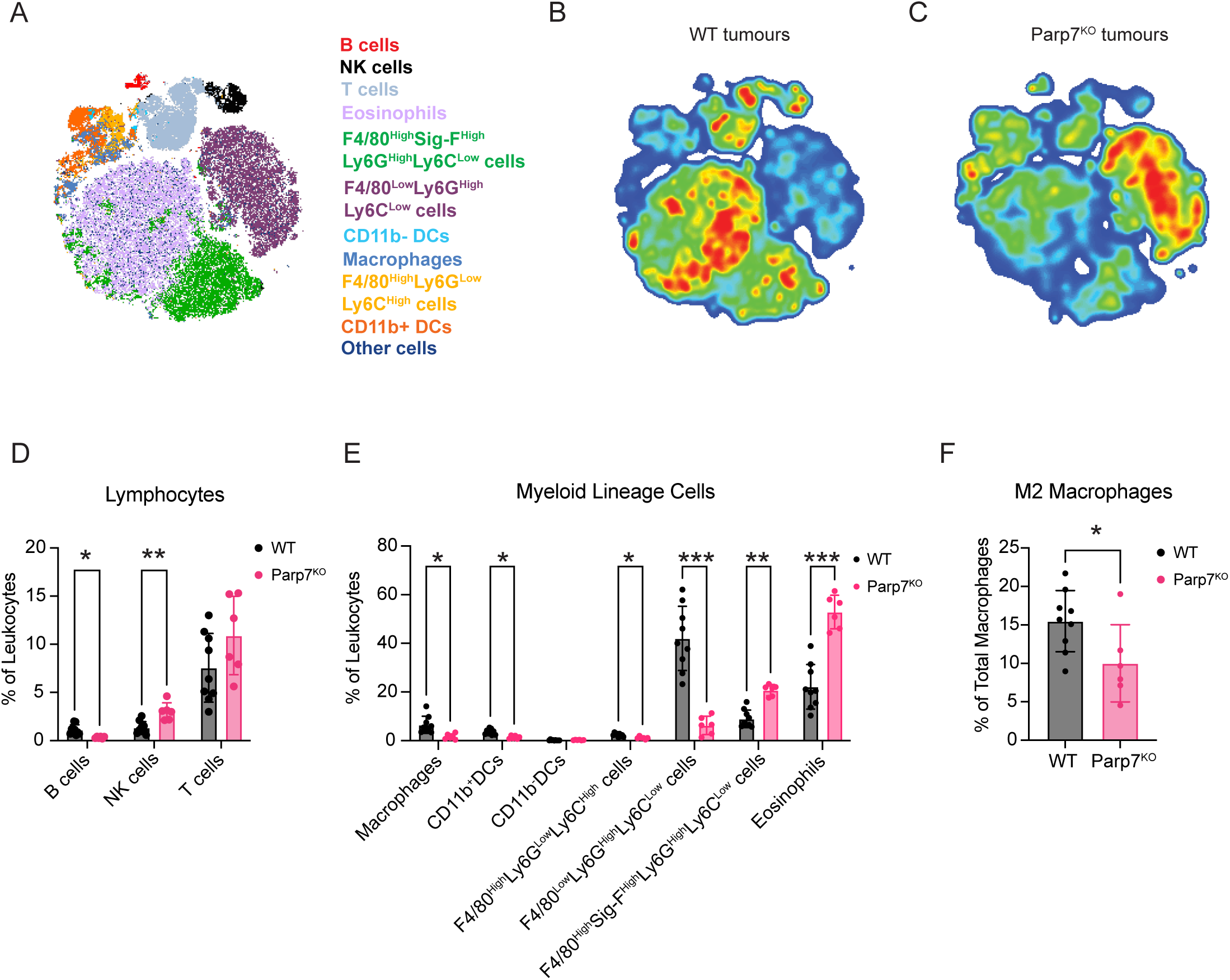

Despite the average proportion of total T cells from Parp7^KO^ tumours being slightly higher than that of WT, this was not statistically significant (**Figure 5D**). To better understand the effect of Parp7 loss on T cell tumour infiltration, we also used a dedicated spectral flow cytometry panel capable of differentiating cancer-relevant T cell subpopulations including cytotoxic (CD8), helper (CD4), and TCRγ8 T cells (**Figure 6A-C; Supplementary Figure S6**). We next determined the proportion of each of these major T cell populations, reported as their abundance relative to the total number of CD4^+^, CD8^+^, and TCRγ8 T cells. A higher proportion of cytotoxic CD8^+^ T cells was observed in Parp7^KO^ tumours relative to CD4 and TCRγ8 T cells (**Figure 6D**). Although the difference in the proportion of CD4^+^ T cells was not statistically significant, Parp7^KO^ tumours with the highest relative CD8^+^ T cell infiltration also had the lowest relative CD4^+^ T cell infiltration (**Figure 6E**). Within the CD4^+^ T cell compartment, there were more T effector/memory (T_EM_; CD44^+^, CD62L^-^) and less T regulatory (Treg; FoxP3^+^) cells in Parp7^KO^ compared with WT tumours (**Figure 6F**). This was specifically reflected in the relative proportion of cells expressing high levels of the immune checkpoint protein PD-1, with no difference in the proportion of PD-1^Low^ CD4^+^ T_EM_ or Tregs. Unlike CD4^+^ T cells, there was no significant difference in the proportion of CD8^+^ T cells that were PD-1^High^ (**Figure 6G**). However, a higher proportion of TCRγ8 T cells were PD-1^High^ in Parp7^KO^ tumours (**Figure 6H**). Overall, these findings show that Parp7^KO^ PDAC tumours had a higher proportion of cytotoxic CD8^+^ T cell infiltration, as well as a lower relative amount of PD-1^High^ CD4^+^ Tregs vs. effector memory CD4^+^ T cells. The higher ratio of effector T cells:Tregs coupled with the lower proportion of M2 macrophages and increased relative amounts of cytotoxic NK cells suggest that loss of *Parp7* may counteract the development of an immunosuppressive TME.

**Figure 6.**
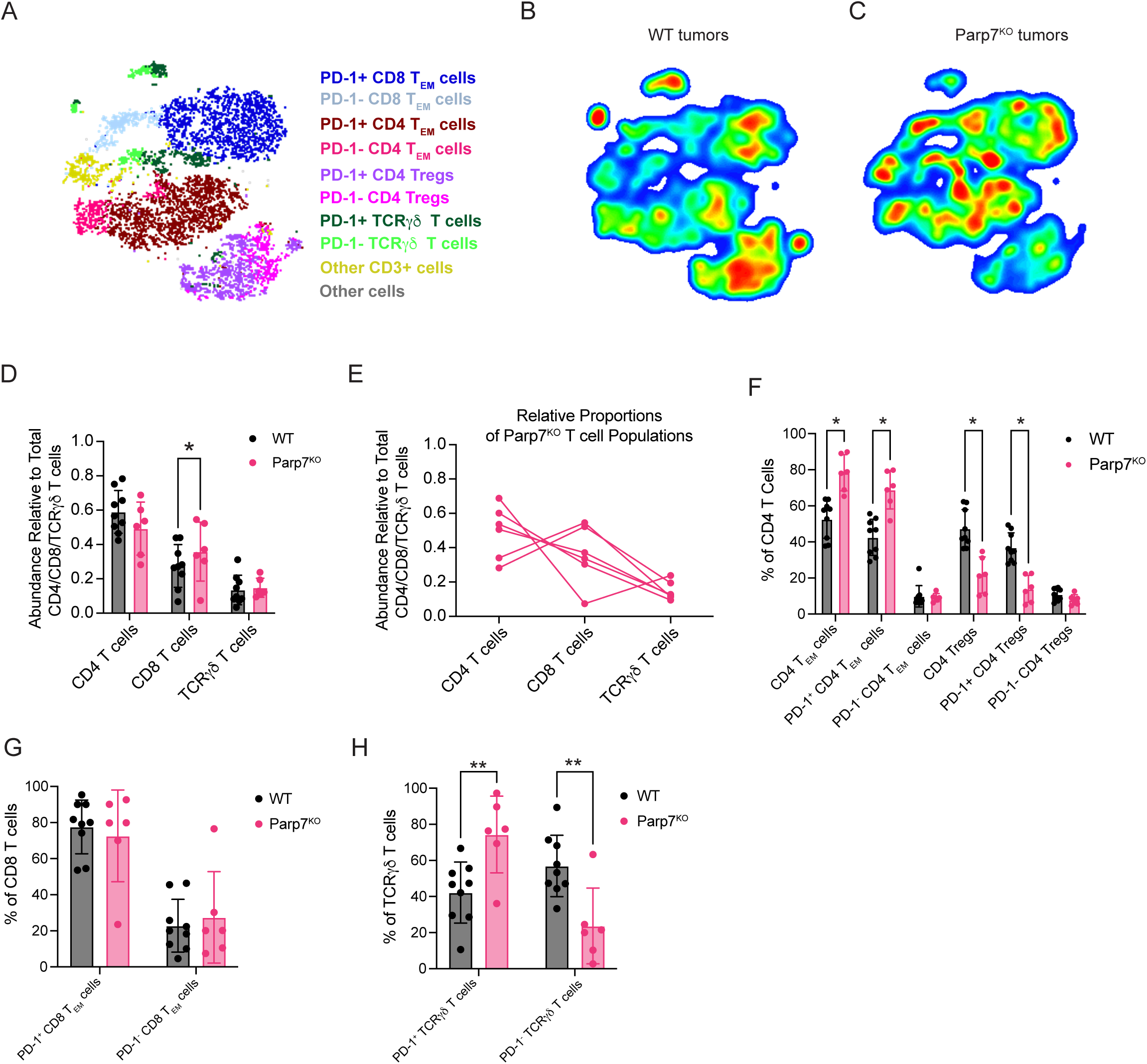

### Gene expression profiling of Parp7^KO^ tumours reveals increased activation of interferon and other immune signalling pathways

RNA sequencing was then used to investigate gene expression profiles and identify cellular pathways that might be differentially regulated between WT and Parp7^KO^ tumours. Principal component analysis revealed distinct clustering between WT and Parp7^KO^ tumours (**Figure 7A**). Hierarchical clustering in the form of a heatmap was used to show distinct expression patterns for overlapping differentially expressed genes (DEGs) between Parp7^KO^ and WT tumours (**Figure 7B**). We identified 1630 significantly changed genes of which 1169 were increased and 461 were decreased in Parp7^KO^ vs WT tumours (**Figure 7C; Supplementary Table S3**). Gene expression profiling confirmed increased expression levels of *Parp9*, *Parp10*, *Parp12* and *Parp14* in Parp7^KO^ tumours. *Parp7* (*Tiparp*) levels were also increased in *Parp7* deficient tumours, which might be due to Parp7 levels in TILs. *Cxcl10* and its receptor, *Cxcr3*, as well as *granzyme A* (*Gzma*) and *granzyme B* (*Gzmb*) levels were elevated in Parp7^KO^ tumours which would contribute to the chemoattraction of immune cells and in killing of cancer cells (**Supplementary Table S3**). Pathway analysis using the Reactome database revealed increases in many immune regulated pathways in Parp7^KO^ tumours including immunoregulatory interactions, interferon signalling pathways, complement activation, neutrophil degranulation, as well as in GPCR signal pathways (**Figure 7D**). These data are consistent with PARP7’s role in regulating interferon signalling and suggest that the loss of PARP7 results in increased inflammation in the TME that may lead to increased immune infiltration and enhanced anti-tumour responses.

**Figure 7.**
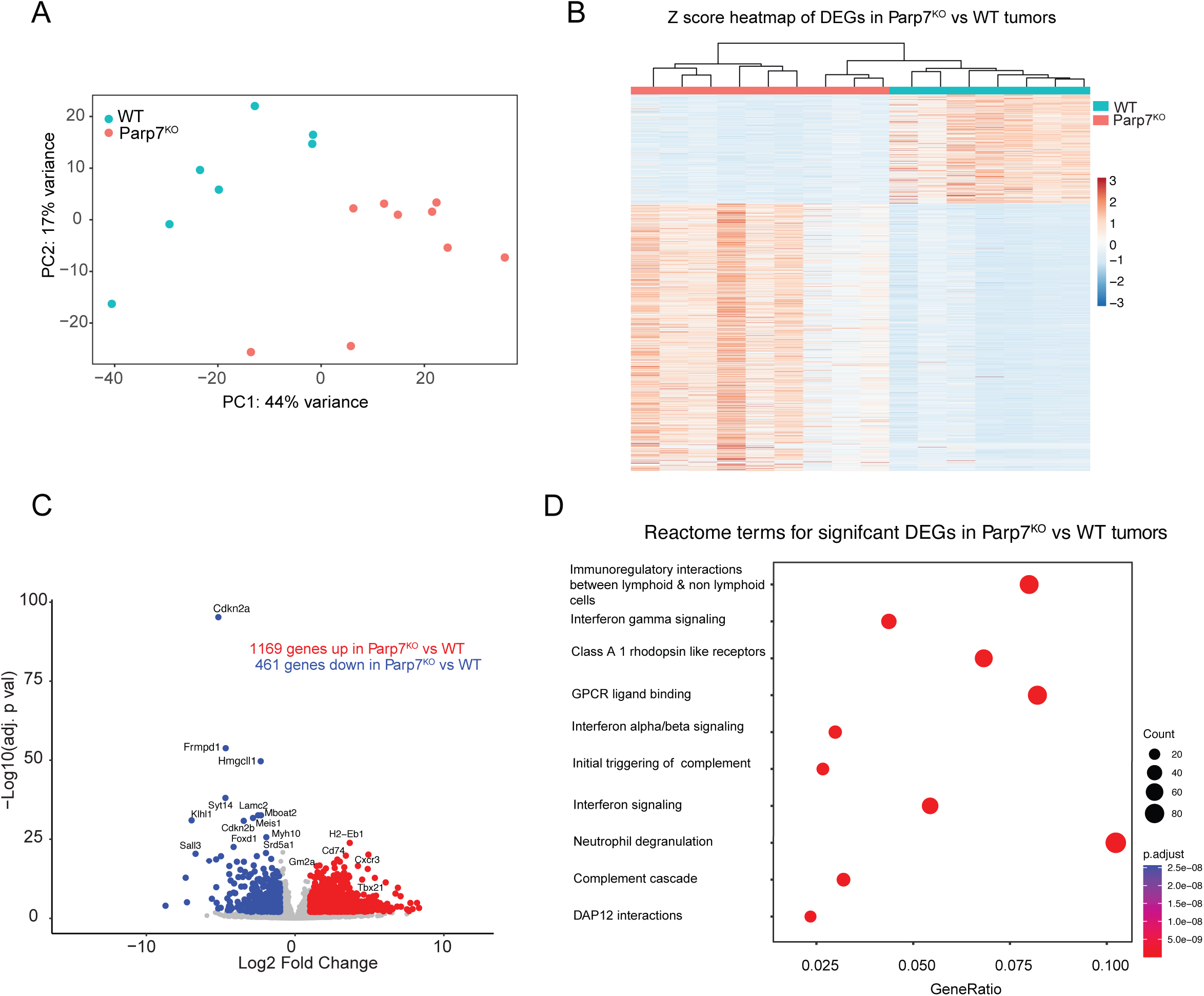

## Discussion

Pharmacological inhibition or loss of PARP7 reduces cancer cell growth in preclinical tumour models of colon and breast cancer through increased immune cell infiltration and antitumor immunity. Antitumor activity following PARP7 inhibition is dependent on CD8 T cells and enhances cancer immunotherapy in combination with anti-PD-1 immune checkpoint inhibition treatment [40]. Since cancer immunotherapy has so far been unsuccessful against PDAC, we were interested to characterize PARP7-dependent IFN-I signalling and tumour growth in a model of PDAC. Using CR705 cells, a mouse PDAC cell line from a spontaneously developed pancreatic tumour in the KPC mouse, we show that loss of *Parp7* increases IFN signalling and reduces tumour growth due to increased immune cell infiltration.

Previous studies have reported that the anti-proliferative effects following PARP7 inhibition depend on multiple factors including AHR [38, 45]. In agreement with these data, CR705 cells express AHR and loss or inhibition of PARP7 reduced their proliferation compared with WT cells. We also observed that AHR signalling was increased in *Parp7* deficient cells, which is consistent with PARP7 acting as a negative regulator of AHR [23, 24]. Despite studies reporting that loss or inhibition of PARP7 enhances AHR signalling, how this crosstalk affects IFN-I signalling remains unclear [23, 37, 38]. Our previous studies in EO771 cells show that PARP7 loss or inhibition increases IFN-I signalling in the absence of AHR [27]. Since IFNβ has been described to increase AHR expression, we cannot exclude that the higher levels of *Ifnb1* contribute to elevated AHR protein levels in Parp7^KO^ cells.

We noted that the expression levels of *Ifnb1* in CR705 cells were lower than those observed in other cell lines, such as EO771, and they did not significantly differ between the WT and Parp7^KO^ cells [27]. Induction of *Ifnb1* was not further increased by treatment with the mouse-specific STING-agonist DMXAA. In line with this, we failed to detect STING expression in CR705 WT or Parp7^KO^ cells. Thus, it would be intriguing to investigate whether the induction or overexpression of STING could enhance the expression of *Ifnb1*, and whether this effect would be amplified in PARP7 deficient cells. Since downstream IFN-I signalling is mediated by ISGF3, we evaluated whether these proteins were affected by *Parp7* loss. In agreement with previous findings, we observed that all three proteins in the ISGF3 complex were upregulated in the Parp7^KO^ cells [27]. However, unlike our previous studies on PARP7 inhibition and knockout in EO771 cells, selected target genes of ISGF3 signalling were upregulated in response to *Parp7* loss but this was not observed in WT cells after PARP7 inhibition. This could be due to adaptation to the loss of PARP7 expression or clonal selective pressures during the isolation of the Parp7^KO^ clone. Despite repeated screening, we were only able to isolate a single Parp7^KO^ clone, so we were unable to confirm these findings in a different knockout clone. This could partly be due to observation that CR705 cells are sensitive to the antiproliferative actions of PARP7 inhibition, making it technically difficult to isolate viable cell clones.

To test downstream IFNAR signalling, we exposed the cells to IFNβ. This resulted in phosphorylation of STAT1 and increased expression levels of target genes, which were substantially higher in the Parp7^KO^ cells. Although the relative levels of pSTAT1 compared to native STAT1 were lower in the Parp7^KO^ cells, the total amount of pSTAT1 was higher. A possible explanation for this could be that increased signalling upregulates negative regulators of the pathway, which function by repressing phosphorylation of STAT1 [46]. Nonetheless, these results imply that loss of PARP7 renders the cells more sensitive to IFN-I signalling. The exact role of ISGF3 signalling in cancer is context specific, as acute signalling is associated with anti-tumour activity, while prolonged signalling promotes tumorigenesis [47]. Previous studies have found that PARP7 represses the IFN-I pathway by MARylating TBK1 [25]. We did not observe any differences between the WT and Parp7^KO^ cells in expression levels of *Ifnb1*, but we observed clear increases in ISGF3 protein levels and downstream IFN-I signalling. This suggests that PARP7 also regulates mechanisms downstream of TBK1, which agrees with a recent study investigating PARP7 signalling in CT26 colon cancer cells [26]. The exact mechanisms behind the elevated ISGF3 protein levels in the absence of PARP7 remain elusive and warrant further investigation to fully understand the function of PARP7 in IFN-I signalling. Consistent with our previous studies in EO771 mouse mammary cells, *Parp7* knockout affected the expression levels of other members of the ARTD family in CR705 cells [27]. We found that *Parp9*, *Parp10* and *Parp14* were upregulated in the CR705 Parp7^KO^ cells. Whether this is due to off-target effects of CRISPR/Cas9, compensation due to the lack of PARP7, or loss of PARP7-dependent regulation of the expression levels of ARTD family members is unclear. PARP9 is a positive regulator of the IFN-I response, and *PARP10* is induced by IFN-Is, and involved in antiviral responses [48, 49]. *PARP14* is regulated by IFN-Is, and in turn acts as a positive regulator of the response [42]. Thus, it is possible that the elevated *Parp14* levels contribute to some of the phenotypes observed in the Parp7^KO^ cells. Furthermore, we observed decreased levels of *Parp2*, *Tnks2* and *Parp13*. TNKS2 and PARP13 are involved in antiviral immunity [50, 51]. It would thus be interesting to examine the effects of knockdown, inhibition or induction of these proteins and assess whether this would affect the phenotypes observed in *Parp7* deficient cells.

Studies in immune incompetent NSG mice, revealed that tumours derived from Parp7^KO^ cells were smaller than those from WT cells. These findings agreed with the differences in cell proliferation observed *in vitro*. However, when experiments were repeated in immunocompetent mice a greater reduction in the growth of Parp7^KO^ compared with WT tumours was observed. This suggests that the immune system more efficiently targets the *Parp7* deficient tumours, which agrees with PARP7 acting as a negatively regulating IFN-I signalling. Gene expression profiling revealed increases in IFN-I and IFNγ signalling as well as other immune signalling pathways in Parp7^KO^ tumours. This increased inflammatory gene profile might also contribute to the increased tumour infiltration of T cells and NK cells as well as reduced levels of anti-inflammatory M2 macrophages which most likely contribute to the increased anti-tumour immunity and reduced tumour progression of Parp7^KO^ tumours. We also observed increased populations of eosinophils in Parp7^KO^ tumours. However, the exact roles of granulocyte populations in cancer are complex, as they can exhibit both pro– and anti-tumorigenic functions that vary by cancer type [52, 53]. Additional studies are needed to determine the important of eosinophils and other granulocytes in PARP7 inhibition-dependent antitumor immunity as well as in PDAC. Importantly, we detected a lower relative abundance of MDSC populations and CD4^+^ Tregs within Parp7^KO^ tumours, further suggesting a shift towards a less immunosuppressive TIL repertoire. Since subcutaneous injection of cancer cells, as was done in this study, does not accurately recapitulate the complex interactions and cell composition of TME that occur in orthotopic injection models or in KPC mouse models, the observed TIL profile is certainly influenced by the method used. It will be necessary to comprehensively investigate the role of PARP7 and its inhibition in more complex PDAC models that better reflect human disease. Collectively, our findings suggest that *Parp7* deficiency in cancer cells increases infiltration of immune cells and enhances anti-tumour activity. The data also suggest that PARP7 may be a suitable target for pancreatic cancer therapy, either alone or in combination with traditional immunotherapy strategies.

## Data availability statement

RNA sequencing data discussed in this paper is available through NCBI’s Gene Expression Omnibus with the accession number GSE276293.

## CRediT authorship contribution statement

Vinicius Kannen: Formal analysis, investigation, methodology, visualization, writing – original draft. Marit Rasmussen: Formal analysis, investigation, visualization, writing – original draft. Siddhartha Das: Formal analysis, visualization. Paolo Giuliana: Formal analysis, investigation, methodology, visualization. Fauzia N. Izzati: Formal analysis, investigation. Hani Choksi: Formal analysis, investigation. Linnea A. M. Erlingsson: Investigation. Ninni E. Olafsen: Investigation. Paola Cappello: Resources. Indrek Teino: Conceptualization, funding acquisition. Toivo Maimets: Conceptualization, funding acquisition. Kristaps Jaudzems: Conceptualization, funding acquisition. Antanas Gulbinas: Conceptualization, funding acquisition. Žilvinas Dambrauskas: Conceptualization, funding acquisition, resources. Resources. Landon Edgar: Conceptualization, supervision. Denis M. Grant: Conceptualization, supervision, project administration. Jason Matthews: Conceptualization, funding acquisition, investigation, supervision, visualization, writing – original draft, project administration. All authors were involved in writing – review & editing.

## Abbreviations

TIPARP: 2,3,7,8-tetrachlorodibenzo-*p*-dioxin (TCDD)-inducible poly-ADP-ribose polymerase
DMXAA: 5,6-dimethylxanthenone-4-acetic acid
ARTD: ADP-ribosyltransferase diphtheria-like
ANOVA: analysis of variance
AHR: aryl hydrocarbon receptor
cGAS: cyclic GMP-AMP synthase
cGAMP: cyclic guanosine monophosphate-adenosine monophosphate
DAMPs: damage associated molecular patterns
DCs: dendritic cells
IRF3: interferon regulatory factor 3
IFNAR: interferon α/β receptor
IRF9: interferon regulatory factor 9
ISGF3: interferon stimulated gene factor 3
KPC: *LSL-Kras^G12D/+^; LSL-Trp53^R172H/+^; Pdx1-Cre*
Gzma: granzyme A
Gzmb: granzyme B
MARylation: mono-ADP-ribosylation
NK: natural killer
PC: pancreatic cancers
PDAC: pancreatic ductal adenocarcinoma
PAMPs: pathogen associated molecular patterns
PRRs: pattern recognition receptors
PD-L1: programmed death ligand 1
PD-1: programmed death receptor
STAT1: signal transducer and activator of transcription 1
STAT2: signal transducer and activator of transcription 2
STING: stimulator of interferon response cGAMP interactor
TBK1: TANK binding kinase 1
TIL: tumour infiltrating leukocyte
TME: tumour microenvironment
IFN-I: type I interferon
WT: wildtype

