## Supplemental Figures 1-6 for "Loss of PARP7 increases type I interferon signalling and prevents pancreatic tumour growth by enhancing immune cell infiltration"

WT Parp7

GAATTCGGCTTTGCAGATTTTTGCATAGCTTTTGAATCTTCATTTCTCAGTTTAAAAAAAGAAAATTGACCTGTA
AGAGCTAACTATAATGCAAGCAGTGATTGCAGATAGTTAACCATAGACTAACGCAAAATGTTTTAATGAATGAAT
GGGTTCAGTTGTCAGTTTTTAAATGATCATCTTTCTTCCTTTCCTGGTAGGATTTGTAGATACTGAGGCACAGTT
GGGAGTTAATCACATC**ATG**GAAGTGGAAACCACTGAACCTGAGCCAGACTGTGTAGTACAGCCTCCTTCTCCTTC
TGATGACTTTTCATGCCA**AATGAGAATTTCTGAGAAGA**TCTCTCCATTGAAAACGTGTTTTAAGAAAAAACAGGA
ACAAAAAAGATTGGGAACTGGAACCCTGAGATCCTTGAGGCCAATATTAAATACTTTGCTAGAATCTGGCTCACT
TGATGGAGTTTTTAGAGCTAGAGACCAAAACAGAGATGAGAGCAGCTTACATGAACATATAGTGAAAAAACCCCT
GGAAATCAACCCATCGTGTCCACCAGCAGAAAACAGTATGCCTGTCCTGATTCCTGATGGGACAAATGTTGAGGG
CCAATTACCAGAAGCGCATCCTTCTACAGATGCTCCAGAACAGGGGGTTCCAATCCAAGACCACAGTTTTCCACC
AGAAACCATCAGTGGGACAGTGGCAGATTCTACAACAGGACACTTCCAAACTGACCTTTTGCATCCTGTTTCAGG
TGATGTTCCTACAAGTCCTGACTGCGTAGATAAAGTTATGGATTATGTACCAGGAGCTTTCCAAGACAAAAGCCG
AATTC

Missing 104 (7/17)

GAATTCGGCTTTGCAGATTTTTGCATAGCTTTTGAATCTTCATTTCTCAGTTTAAAAAAAGAAAATTGACCTGTA
AGAGCTAACTATAATGCAAGCAGTGATTGCAGATAGTTAACCATAGACTAACGCAAAATGTTTTAATGAATGAAT
GGGTTCAGTTGTCAGTTTTTAAATGATCATCTTTCTTCCTTTCCTGGTAGGATTTGTAGATACTGAGGCACAGTT
GGG**AGAAGA**TCTCTCCATTGAAAACGTGTTTTAAGAAAAAACAGGAACAAAAAAGATTGGGAACTGGAACCCTGA
GATCCTTGAGGCCAATATTAAATACTTTGCTAGAATCTGGCTCACTTGATGGAGTTTTTAGAGCTAGAGACCAAA
ACAGAGATGAGAGCAGCTTACATGAACATATAGTGAAAAAACCCCTGGAAATCAACCCATCGTGTCCACCAGCAG
AAAACAGTATGCCTGTCCTGATTCCTGATGGGACAAATGTTGAGGGCCAATTACCAGAAGCGCATCCTTCTACAG
ATGCTCCAGAACAGGGGGTTCCAATCCAAGACCACAGTTTTCCACCAGAAACCATCAGTGGGACAGTGGCAGATT
CTACAACAGGACACTTCCAAACTGACCTTTTGCATCCTGTTTCAGGTGATGTTCCTACAAGTCCTGACTGCGTAG
ATAAAGTTATGGATTATGTACCAGGAGCTTTCCAAGACAAAAGCCGAATTC

Missing 31 (10 of 17)

GAATTCGGCTTTGCAGATTTTTGCATAGCTTTTGAATCTTCATTTCTCAGTTTAAAAAAAGAAAATTGACCTGTA
AGAGCTAACTATAATGCAAGCAGTGATTGCAGATAGTTAACCATAGACTAACGCAAAATGTTTTAATGAATGAAT
GGGTTCAGTTGTCAGTTTTTAAATGATCATCTTTCTTCCTTTCCTGGTAGGATTTGTAGATACTGAGGCACAGTT
GGGAGTTAATCACATC**ATG**GAAGTGGAAACCACTGAACCTGAGCCAGACTGTGTAGTACAGCCTCCTTCTCCTTC
TGATGACTCTCTCCATTGAAAACGTGTTTTAAGAAAAAACAGGAACAAAAAAGATTGGGAACTGGAACCCTGAGA
TCCTTGAGGCCAATATTAAATACTTTGCTAGAATCTGGCTCACTTGATGGAGTTTTTAGAGCTAGAGACCAAAAC
AGAGATGAGAGCAGCTTACATGAACATATAGTGAAAAAACCCCTGGAAATCAACCCATCGTGTCCACCAGCAGAA
AACAGTATGCCTGTCCTGATTCCTGATGGGACAAATGTTGAGGGCCAATTACCAGAAGCGCATCCTTCTACAGAT
GCTCCAGAACAGGGGGTTCCAATCCAAGACCACAGTTTTCCACCAGAAACCATCAGTGGGACAGTGGCAGATTCT
ACAACAGGACACTTCCAAACTGACCTTTTGCATCCTGTTTCAGGTGATGTTCCTACAAGTCCTGACTGCGTAGAT
AAAGTTATGGATTATGTACCAGGAGCTTTCCAAGACAAAAGCCGAATTC

**Supplementary Figure S1.** Indels resulting in reading frame errors in the Parp7 gene. Genomic DNA was isolated from multiple selected and expanded clones, and the region surrounding the gRNA target site in *Parp7* (bold red font) was amplified and sequenced. The sequences above show the deletions (and frequencies) from the $Parp7^{KO}$ clone used in the study. The start ATG is in bold font. The two nucleotides flanking the deleting sequences are underlined.

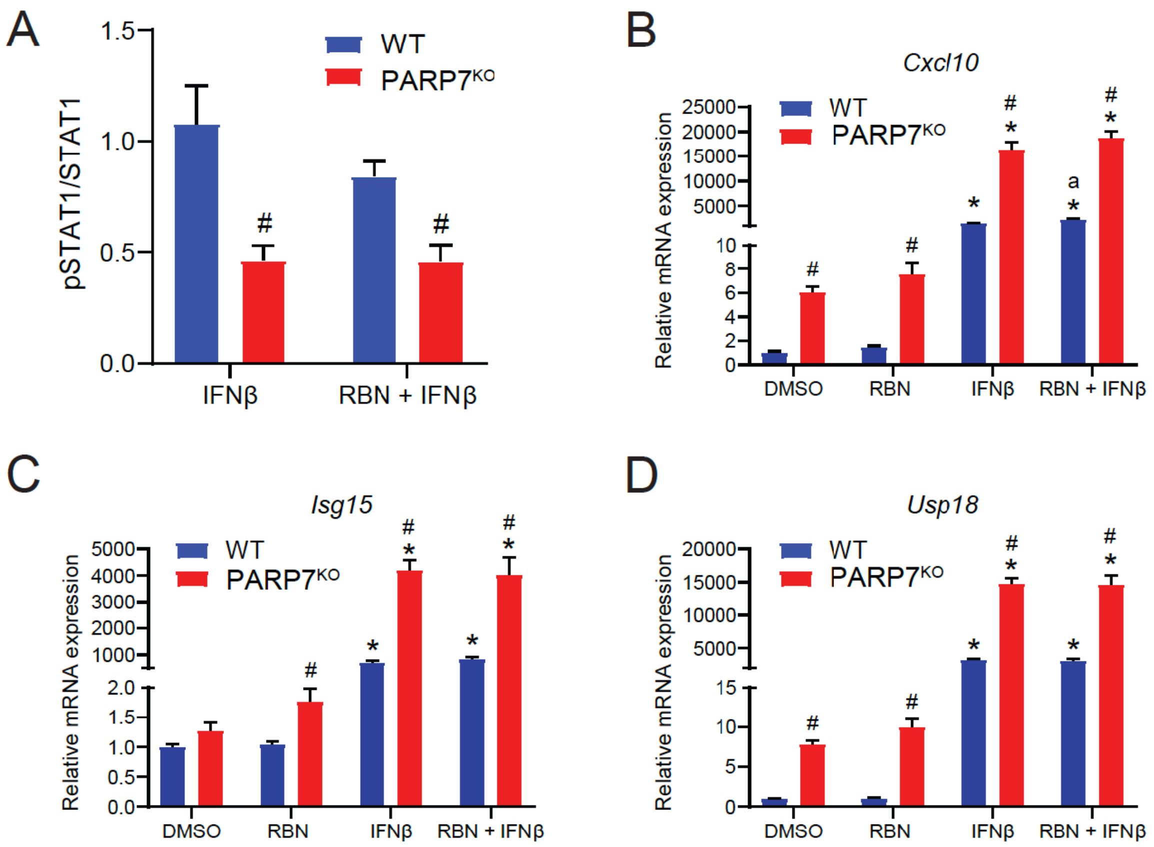


**Supplementary Figure S2.** Responses to exogenous IFNβ. **(A)** Relative pSTAT1 levels. Quantification of pSTAT1 bands relative to native STAT1 bands (pSTAT1/STAT1). Cells were pre-treated with 100 nM of RBN-2397 for 24 h and exposed to 1000 U/mL of IFNβ for 1 h. **(B-D)** Split axis showing increased basal expression levels of *Cxcl10*, *Isg15* and *Usp18* in Parp7$^{KO}$ cells. Cells were pre-treated with RBN-2397 for 24 h and exposed to 1000 U/mL of IFNβ for 4 h. *$p$<0.05 denotes statistical significance compared with DMSO treated samples, #$p$<0.05denotes significance due to loss of PARP7.

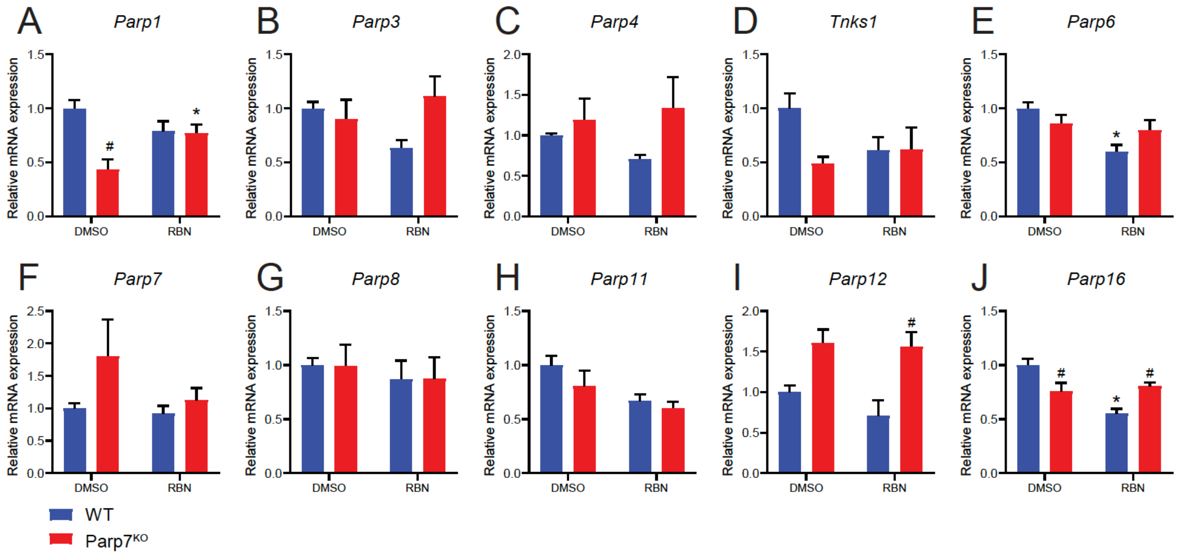


**Supplementary Figure S3.** Expression levels of ARTD family members in CR705 WT and Parp7$^{KO}$ cells after treatment with RBN-2397. (A) Levels of *Parp1* were decreased in the DMSO-treated knockout cells. (B-D, F-H) Levels of *Parp3*, *Parp4*, *Tnks1*, *Parp6*, *Parp7*, *Parp8* and *Parp11* were not significantly affected by loss or inhibition of PARP7. (E) *Parp6* expression was lower in the WT cells treated with RBN-2397. (J) *Parp12* levels were significantly elevated in the Parp7$^{KO}$ cells after treatment with RBN-2397. (J) Levels of *Parp16* were significantly lower in the DMSO treated Parp7$^{KO}$ cells, and in WT cells treated with RBN-2397. Cells were treated with DMSO or 100 nM of RBN-2397 for 24 h, and expression levels were determined with RT-qPCR. *$p<0.05$ denotes statistical significance from the DMSO treated samples, #$p<0.05$ significance due to loss of PARP7.

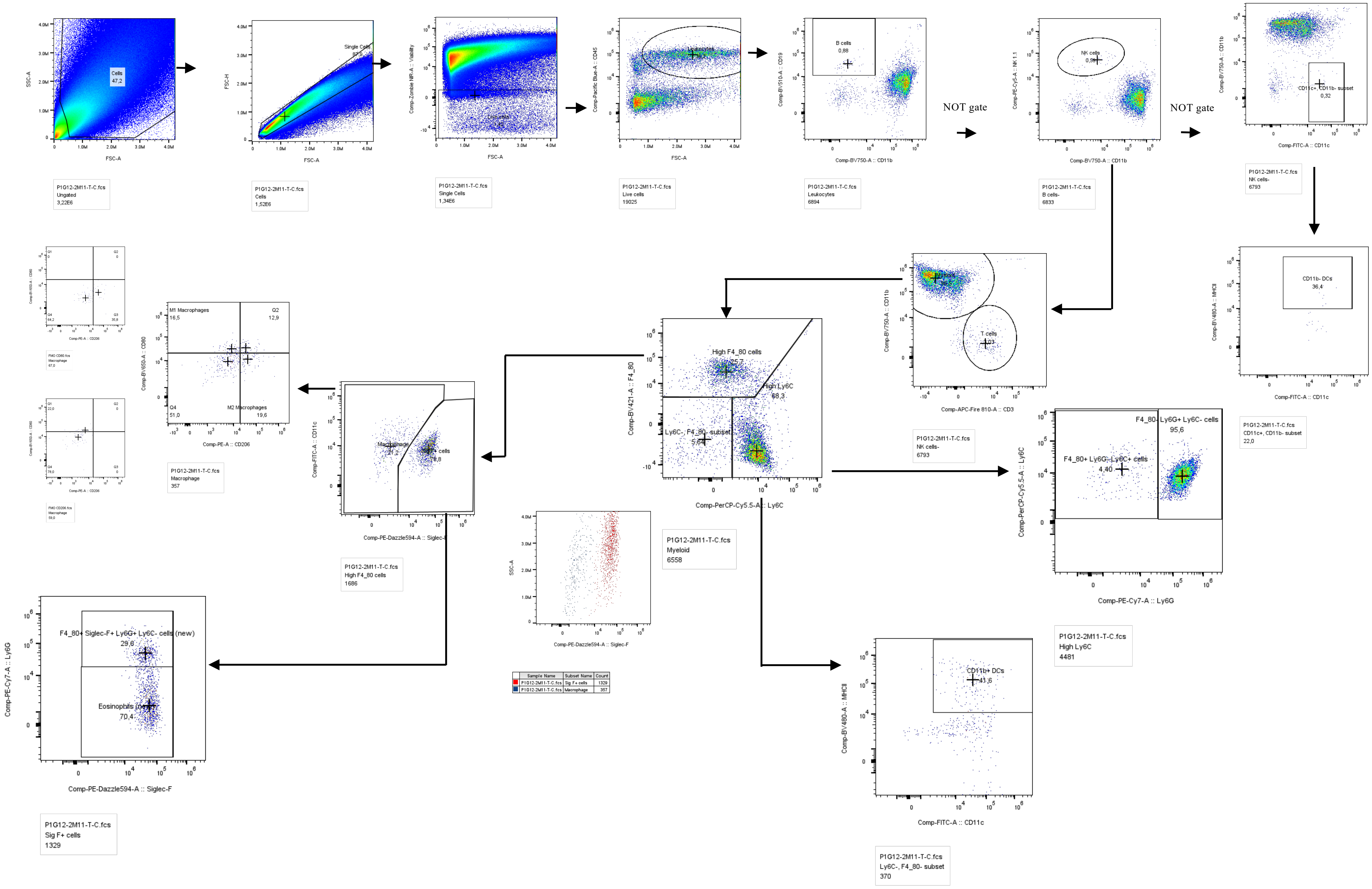


**Supplementary Figure S4.** Gating strategy for flow cytometry analysis of leukocytes from single cell suspensions of WT and Parp7$^{KO}$ tumours.

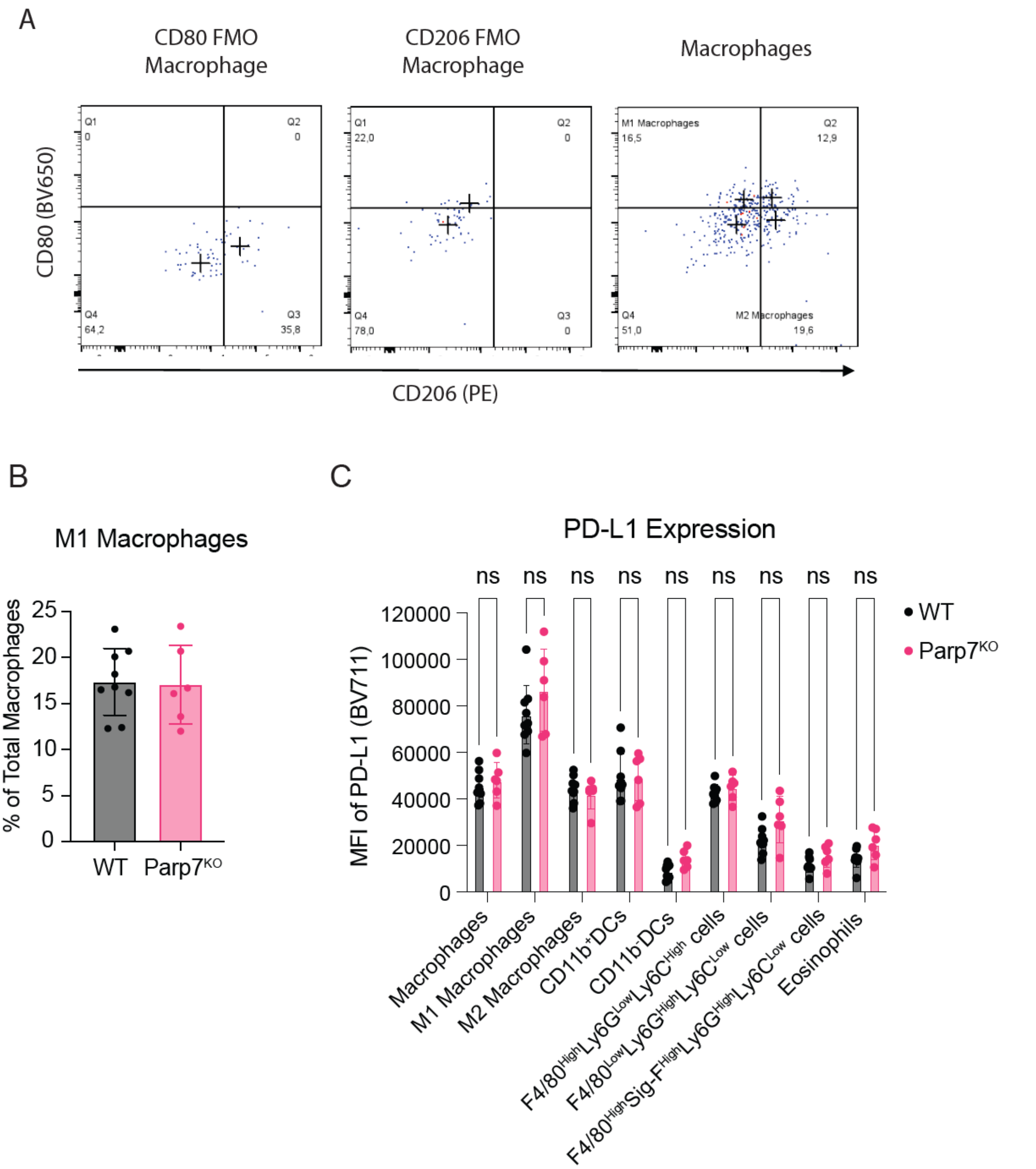


**Supplementary Figure S5.** Flow cytometry analysis of macrophage phenotypes in Parp7$^{KO}$ and WT tumours. **(A)** Analysis of M1-like (CD80$^{hi}$, CD206$^{low}$) pro-inflammatory or M2-like (CD80$^{low}$, CD206$^{hi}$) anti-inflammatory phenotype and macrophages in tumours. **(B)** M1 macrophages from Parp7$^{KO}$ and WT tumours. **(C)** No differences in the expression of the immune checkpoint PD-L1 were observed across all tumour infiltrating leukocyte (TIL) populations between Parp7$^{KO}$ and WT tumours.

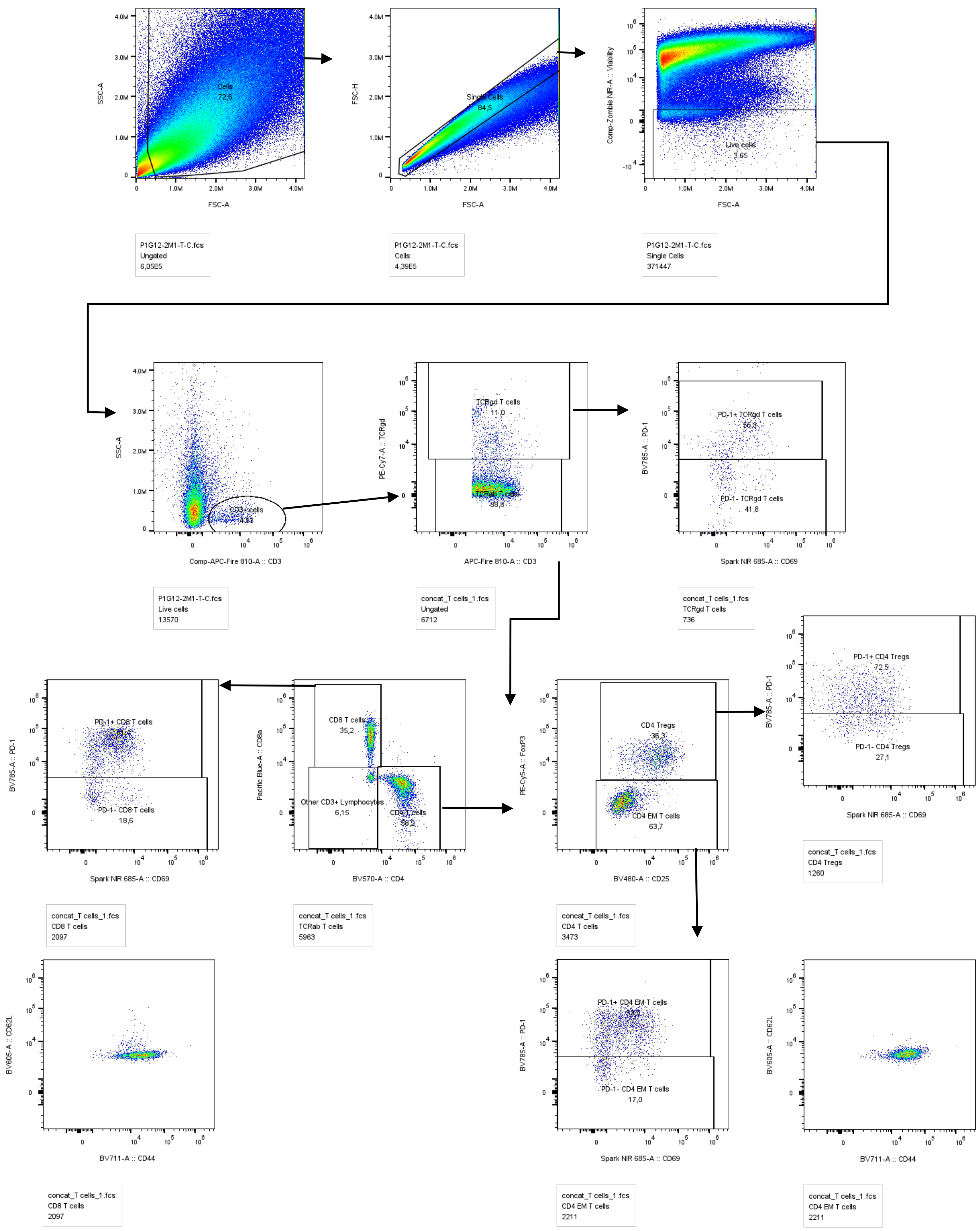


**Supplementary Figure S6.** Gating strategy for flow cytometry analysis of T cell populations from single cell suspensions of WT and Parp7$^{KO}$ tumours.
