## Supplemental Table 1 for "Loss of PARP7 increases type I interferon signalling and prevents pancreatic tumour growth by enhancing immune cell infiltration"

**Supplementary Table S1.** Primers used for RT-qPCR.

| *Gene* | Strand | Sequence |
|---|---|---|
| *Tbp* | Forward | GCACAGGAGCCAAGAGTGAA |
| | Reverse | TAGCTGGGAAGCCCAACTTC |
| *Ifnb1* | Forward | TGGGAGATGTCCTCAACTGC |
| | Reverse | CCAGGAGTAGCTGTTGTACT |
| *Cyp1a1* | Forward | CGTTATGACCATGATGACCAAGA |
| | Reverse | TCCCCAAACTCATTGCTCAGAT |
| *Cxcl10* | Forward | CCAAGTGCTGCCGTCATTTTC |
| | Reverse | GGCTCGCAGGGATGATTTCAA |
| *Stat1* | Forward | GCCTCTCATTGTCACCGAAGAAC |
| | Reverse | TGGCTGACGTTGGAGATCACCA |
| *Stat2* | Forward | GAACCAACTCTCCATTGCCTGG |
| | Reverse | CGTAAGAGGAGAACTGCCAGCT |
| *Irf9* | Forward | CAACATAGGCGGTGGTGGCAAT |
| | Reverse | GTTGATGCTCCAGGAACACTGG |
| *Isg15* | Forward | CATCCTGGTGAGGAACGAAAGG |
| | Reverse | CTCAGCCAGAACTGGTCTTCGT |
| *Usp18* | Forward | GGAACCTGACTAAGGACCAGATC |
| | Reverse | GAGAGTGTGAGCAGTTTGCTCC |
| *Parp1* | Forward | GGCAGCCTGATGTTGAGGT |
| | Reverse | GCGTACTCCGCTAAAAAGTCAC |
| *Parp2* | Forward | TGGAAGGCGAGTGCTAAATG |
| | Reverse | GGGCTTTGCCCTTTAACAGC |
| *Parp3* | Forward | TGCGGCATGTTTGGAAAGTG |
| | Reverse | GTGCATGGTGGTAACATAGCC |
| *Parp4* | Forward | AGTGCTACAGCCCGTTTCC |
| | Reverse | CACAGCTTTCAGTTGTGGGC |
| *Tnks1* | Forward | CCCTGAGGCCTTACCTACCT |
| | Reverse | TCAAGACCCGCAACTTCTCC |
| *Tnks2* | Forward | TGATGGCAGAAAGTCAACTCCA |
| | Reverse | GCCACAGGTCCATTGCATTC |
| *Parp6* | Forward | GTACCTTGATGGACCAGAGCC |
| | Reverse | GCCAGCTCGGAACTTCTTGA |
| *Parp7* | Forward | AAAACCCCTGGAAATCAACC |
| | Reverse | GAATCTGCCACTGTCCCACT |
| *Parp8* | Forward | CACTTCCGAAACCACTTCGC |
| | Reverse | TAGGATACACTTTTGGGGCCG |
| *Parp9* | Forward | GCATTTGCTAAAGAGCACAAGGA |
| | Reverse | AAAGCACCACTATTACCGCTGA |
| *Parp10* | Forward | CGAAACGGCACACTCTACGG |
| | Reverse | GAGACCCTCAAAGGAGGTGC |
| *Parp11* | Forward | CAAACCCTTGTGGCTCCATTTCC |
| | Reverse | AGGCACTGATGGAGAAAGGAGC |
| *Parp12* | Forward | AAGTTCTGACCTGGTGAGCAGG |
| | Reverse | TGGTGACACAGGACCTGAACTG |
| *Parp13* | Forward | AGTAGTCCCACTGGTTTTGGC |
| | Reverse | TGCAACTCTGTGGCTTGTGG |
| *Parp14* | Forward | TGCTGAAGCTGTCAAGACTACA |
| | Reverse | ACAATGGCATGGGTCGTAGC |
