## Supplemental Table 2 for "Loss of PARP7 increases type I interferon signalling and prevents pancreatic tumour growth by enhancing immune cell infiltration"

**Supplementary Table S2.** Antibodies and viability dye used in spectral flow cytometry.

| Target | Clone | Conjugate | Vendor | Product code |
|---|---|---|---|---|
| F4/80 | T45-2342 | BV421 | BD Biosciences | 565411 |
| CD45 | 30-F11 | Pacific Blue | BioLegend | 103126 |
| CD8a | 53-6.7 | Pacific Blue | BioLegend | 100725 |
| CD25 | PC61 | BV480 | BD Biosciences | 566202 |
| I-A/I-E (MHC II) | M5/114.15.2 | BV480 | BD Biosciences | 566086 |
| CD19 | 6D5 | BV510 | BioLegend | 115545 |
| CD4 | RM4-5 | BV570 | BioLegend | 100541 |
| CD62L | MEL-14 | BV605 | BioLegend | 104438 |
| CD183 (CXCR3) | CXCR3-173 | BV650 | BioLegend | 126531 |
| CD80 | 16-10A1 | BV650 | BD Biosciences | 563687 |
| CD44 | IM7 | BV711 | BD Biosciences | 563971 |
| PD-L1 | 10F.9G2 | BV711 | Biolegend | 124319 |
| CD11b | M1/70 | BV750 | BioLegend | 101267 |
| CD279 (PD-1) | 29F.1A12 | BV785 | BioLegend | 135225 |
| CD11c | N418 | FITC | BioLegend | 117305 |
| CD206 | C068C2 | PE | BioLegend | 141705 |
| RORy(t) | B2D | PE | Invitrogen | 141705 |
| CD170 (Siglec-F) | S17007L | PE/Dazzle 594 | BioLegend | 155529 |
| FoxP3 | FJK-16s | PE/Cy5 | Invitrogen | 141705 |
| NK1.1 | PK136 | PE/Cy5 | BioLegend | 108715 |
| Ly-6C | HK1.4 | PerCP/Cy5.5 | BioLegend | 128011 |
| Ly-6G | 1A8 | PE/Cy7 | BioLegend | 127617 |
| TCR gamma/delta | GL3 | PE/Cy7 | BioLegend | 118123 |
| CCR6 | 29-2L17 | APC | BioLegend | 129813 |
| CD163 | S15049I | APC | BioLegend | 155305 |
| CD69 | H1.2F3 | Spark NIR 685 | BioLegend | 104557 |
| CD38 | 90 | AF700 | BioLegend | 102741 |
| CD19 | 6D5 | AF700 | BioLegend | 115528 |
| CD3 | 17A2 | APC/Fire 810 | BioLegend | 100267 |
| CD16/32 | 93 | Unconjugated | BioLegend | 101320 |
| Viability Dye | | | | |
| Zombie NIR | n/a | n/a | BioLegend | 423105 |
